# Plasma-derived Extracellular Vesicles (EVs) as Biomarkers of Sepsis in Burn Patients via Label-free Raman Spectroscopy

**DOI:** 10.1101/2024.05.14.593634

**Authors:** Hannah J. O’Toole, Neona Lowe, Vishalakshi Arun, Anna V. Kolesov, Tina L. Palmieri, Nam K. Tran, Randy P. Carney

**Affiliations:** Department of Biomedical Engineering, University of California, Davis, 1 Shields Ave, Davis., CA 95616, USA; Department of Neurobiology, Physiology, and Behavior, University of California, Davis, 1 Shields Ave., Davis, CA 95616, USA; Division of Burn Surgery & Reconstruction, Department of Surgery, University of California, Davis Health, Firefighters Burn Institute Regional Burn Center, 2315 X Street, Sacramento, CA 95616, USA; Shriners Hospitals for Children Northern California, 2425 Stockton Blvd., Sacramento, CA 95817, USA; Department of Pathology and Laboratory Medicine, University of California, Davis, 4400 V. St., Sacramento, CA 95817, USA

**Keywords:** exosomes, diagnostics, systemic inflammatory response syndrome (SIRS), bacterial EVs (bEVs)

## Abstract

Sepsis following burn trauma is a global complication with high mortality, with ∼60% of burn patient deaths resulting from infectious complications. Sepsis diagnosis is complicated by confounding clinical manifestations of the burn injury, and current biomarkers markers lack the sensitivity and specificity required for prompt treatment. Circulating extracellular vesicles (EVs) from patient liquid biopsy as biomarkers of sepsis due to their release by pathogens from bacterial biofilms and roles in subsequent immune response. This study applies Raman spectroscopy to patient plasma derived EVs for rapid, sensitive, and specific detection of sepsis in burn patients, achieving 97.5% sensitivity and 90.0% specificity. Furthermore, spectral differences between septic and non-septic burn patient EVs could be traced to specific glycoconjugates of bacterial strains associated with sepsis morbidity. This work illustrates the potential application of EVs as biomarkers in clinical burn trauma care, and establishes Raman analysis as a fast, label-free method to specifically identify features of bacterial EVs relevant to infection amongst the host background.

---

Sepsis is characterized as a dysregulated systemic response to infection causing life threatening organ dysfunction and devastating consequences if not promptly recognized and treated.^[1]^ Sepsis is a global health crisis accounting for an estimated 19.7% of all global deaths in 2017.^[2]^ Clinical indications of sepsis include an increase in the Sequential (Sepsis-related) Organ Failure Assessment (SOFA) score by 2 points (or about 10% greater chance of in-hospital morbidity). A quick SOFA (qSOFA) may be performed to rapidly identify patients at risk of sepsis and subsequent septic shock.^[3,4]^ This includes presenting with at least two of the following: systolic blood pressure 100mmHg, respiratory rate 22 breaths/min, or altered mental status.^[1]^ Due to the morbidity of organ dysfunction, any patient that presents with infection should be monitored for sepsis. However, due to the heterogeneity of patient and pathogen factors, sepsis can present differently from patient to patient, making it difficult to definitively diagnose.

The clinical phenotype of sepsis becomes further altered when the patient sustains injury due to a temperature related event, e.g., a burn patient. Burn patients have higher susceptibility to bacterial invasion and sepsis due to the compromised skin barrier and already present systemic dysregulation and immunosuppression in response to burn trauma.^[5]^ Burn wound infection is a major cause of burn patient death after the first 24-hours post trauma and resuscitation.^[6]^ Furthermore, diagnosis of sepsis in burn patients is complicated due to confounding clinical manifestations, negating the use of qSOFA and many other traditional criteria for sepsis. Prognosis for septic burn patients is poor, with recent estimates of >60% of burn patient deaths resulting from infectious complications.^[7]^ Before the updated 2016 Journal of the American Medical Association (JAMA) nomenclature for sepsis (known as Sepsis-3)^[1]^, the pathobiology was identified by the presence of two Systemic Inflammatory Response Syndrome (SIRS) criteria. However, these criteria are confounding with the clinical manifestations of severe burns (e.g., higher white blood cell counts, temperature, and changes in heart rate and respiratory rate), among other clinical presentations of inflammation and host response such as compensatory anti-inflammatory response syndrome (CARS), and do not necessarily indicate presence of infection.^[8]^

Several different definition-based diagnostic criteria formulations have been considered for burn patients to predict sepsis (ABA-criteria, BURN-6, FF4, Sepsis-3), however none have both high sensitivity and specificity.^[9–11]^ Investigations into clinical biomarkers of sepsis in burn patients such as cytokine and chemokine markers (interleukins (IL)-6, IL-8, IL-10), metabolites (L-lactate, procalcitonin (PCT)), soluble receptors (HLA-DR, nCD-64), and vascular endothelial cell or coagulation cascade-related markers are under investigation, but with limited viability, especially for early recognition of sepsis in this patient population.^[12–19]^ There is yet to be gold standard diagnostic criteria for burn sepsis identification.^[20]^

Due to the heterogeneity of sepsis presentation in patients, especially in burn patients, individualized dosing of antimicrobial medication is highly important. The time-to-antibiotic treatment is imperative, with an observational study showing a 1.5% increase in mortality per hour delay of antibiotic treatment of patients in an ICU setting.^[21,22]^ The timing of blood cultures for bacteremia have a significant impact on correlation with sepsis, with positive cultures drawn at a median of 4 days after burn center admission in a retrospective review of 282 blood cultures form 196 burn ICU patients.^[21]^ Technical approaches for characterization of monocyte volume in response to pro-inflammatory signals from bacteria have also not been successful in reducing lead time in sepsis detection. Thus, there is a major unmet clinical need for early-stage determination of sepsis in burn patients to ensure adequate treatment and potentially significantly improve patient outcomes.

There is a strong rationale to assess circulating extracellular vesicles (EVs) as biomarkers for sepsis due to their primary release by the pathogens involved in sepsis bacteremia, as well as their many roles in the subsequent inflammatory response, including cell-to-cell communication. EVs are ubiquitous lipid membrane bound nanovesicles that are secreted by all cells across all kingdoms (mammalian, plant, and bacterial) into the extracellular space.^[23]^ EVs are of particular interest because their membrane-bound and internal biomolecular contents (e.g., proteins, lipids, metabolites, genetic material, cytokines) are variable in response to cellular stressors and disease pathogenesis, and can therefore be used as indicators of disease, in cancer, wound healing, neurodegenerative disorders, cardiovascular disease, and more.^[24–28]^ While EVs are central components in key cellular processes, such as contributing to homeostasis in both basic cell functions and immune response,^[29]^ intense research is underway to establish EVs as biomarkers of aberrant cell signaling and pathogenesis. EVs are widely regarded as potent candidates for liquid biopsy-based cancer detection.^[25,30]^ Likewise, the use of EVs for acute injury detection is growing with potential applications cited for sepsis, making EV diagnostics for early detection of sepsis a viable option for routine patient care.^[31]^

In prokaryotes, EVs were originally thought to be produced by gram-negative bacteria, so-called outer membrane vesicles (OMVs), then 30 years after their initial discovery reports of EVs produced by gram-positive bacteria surfaced.^[32–34]^ Thus, identification of EVs released by microbial cells driving infection in burn patient populations is of particular interest, since they may be released in high numbers into circulating biofluids *via* damaged endothelium at the site of burn injury (**Figure 1a**).

**Figure 1:**
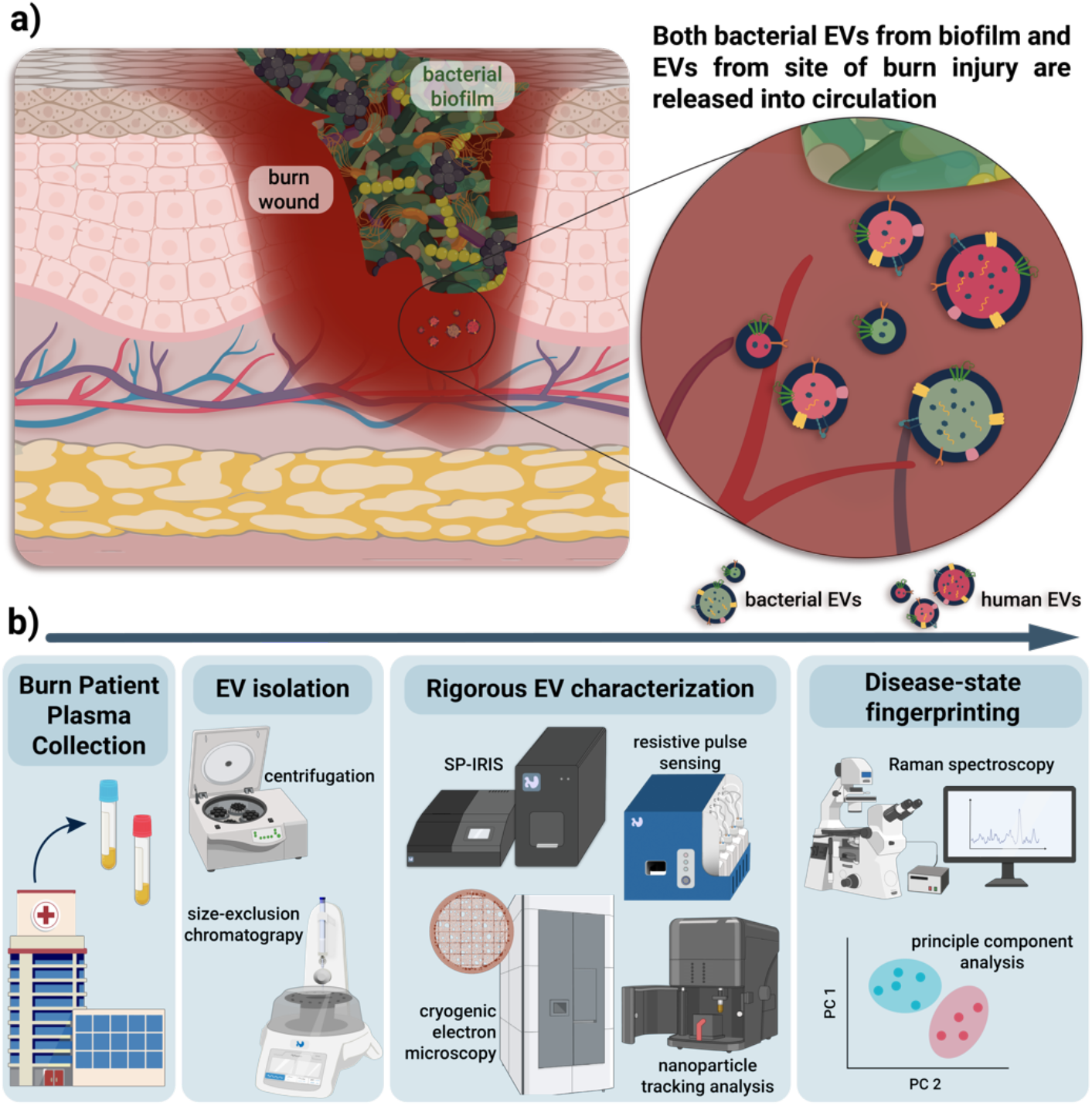
Extracellular vesicle release during bacterial infection in burn injury; Overview schema of sample preparation and acquisition. (a) The rationale behind our work is that EVs originating from colonizing bacteria are released into circulatory biofluids via damaged endothelium at the site of burn injury. Inflammatory response to injury results in secondary release of EVs from immune and endothelial cells as well. (b) Collection and analysis of EVs in circulating biofluids are a way to determine sepsis in advance of currently available methods.

Here we apply label-free Raman spectroscopy (RS) to analyze circulating EVs isolated from patient plasma. RS is a non-destructive, label-free optical spectroscopic method that captures a comprehensive snapshot of the chemical composition of a given biofluid, whereby the signal reflects the complex mixture of lipid, metabolites, proteins, and more in each sample based on changes in their molecular vibrations.^[35–37]^ Chemical spectroscopic analysis of EVs using RS has recently shown great promise as a label-free tool with high sensitivity to detect and monitor disease in a variety of contexts, particularly in liquid biopsy analysis.^[38]^ Several studies have investigated the application of RS to distinguish bacterial EVs released from different pathogen sources.^[39–41]^ We utilize this approach to stablish a comprehensive workflow to evaluate EV liquid biopsy isolated from burn patient plasma samples as a proof-of-concept for sensitive and rapid detection of sepsis (**Figure 1b**).

## Results and Discussion

### Clinical Biofluid Collection

De-identified remnant clinical chemistry lithium heparin plasma samples were collected from adult burn patients from the University of California, Davis Health Fire Fighters Burn Institute Regional Burn Center. Samples utilized the UC Davis Pathology Biorepository anonymous clinical research specimen (ARCS) protocol (IRB#1911992).

In this study, patient plasma from both septic and non-septic burn patients was analyzed (n=14). At the time of collection, two of these patients had not yet been confirmed as septic, but during the course of our study their prognosis was updated, and they were included within the septic burn patient sample cohort (n=8) for analysis. Available de-identified patient information is summarized in **Supplemental Table 1**, including age, sex, procalcitonin level, and total body surface area of burn.

### EV Isolation and Characterization

We first isolated and characterized EVs from patient plasma samples (n=14). Patient plasma was centrifuged to remove cell debris. Size exclusion chromatography (SEC) was then used to isolate EVs from septic (n=8) and non-septic (n=6) (control) burn patient plasma samples. EVs were subsequently characterized by morphology and size, following suggestions from the Minimal Information for Studying Extracellular Vesicles (MISEV) 2023 guidelines and technique specific recommendations for EV characterization as shown in **Figure 2**.^[42,43]^ As suggested by MISEV 2023, EV morphology was further characterized with cryogenic electron microscopy (CryoEM) as a complementary method. CryoEM images of the EVs reveal the expected lipid bilayer structure for both septic and non-septic samples (**Figure 2a–d**).

**Figure 2:**
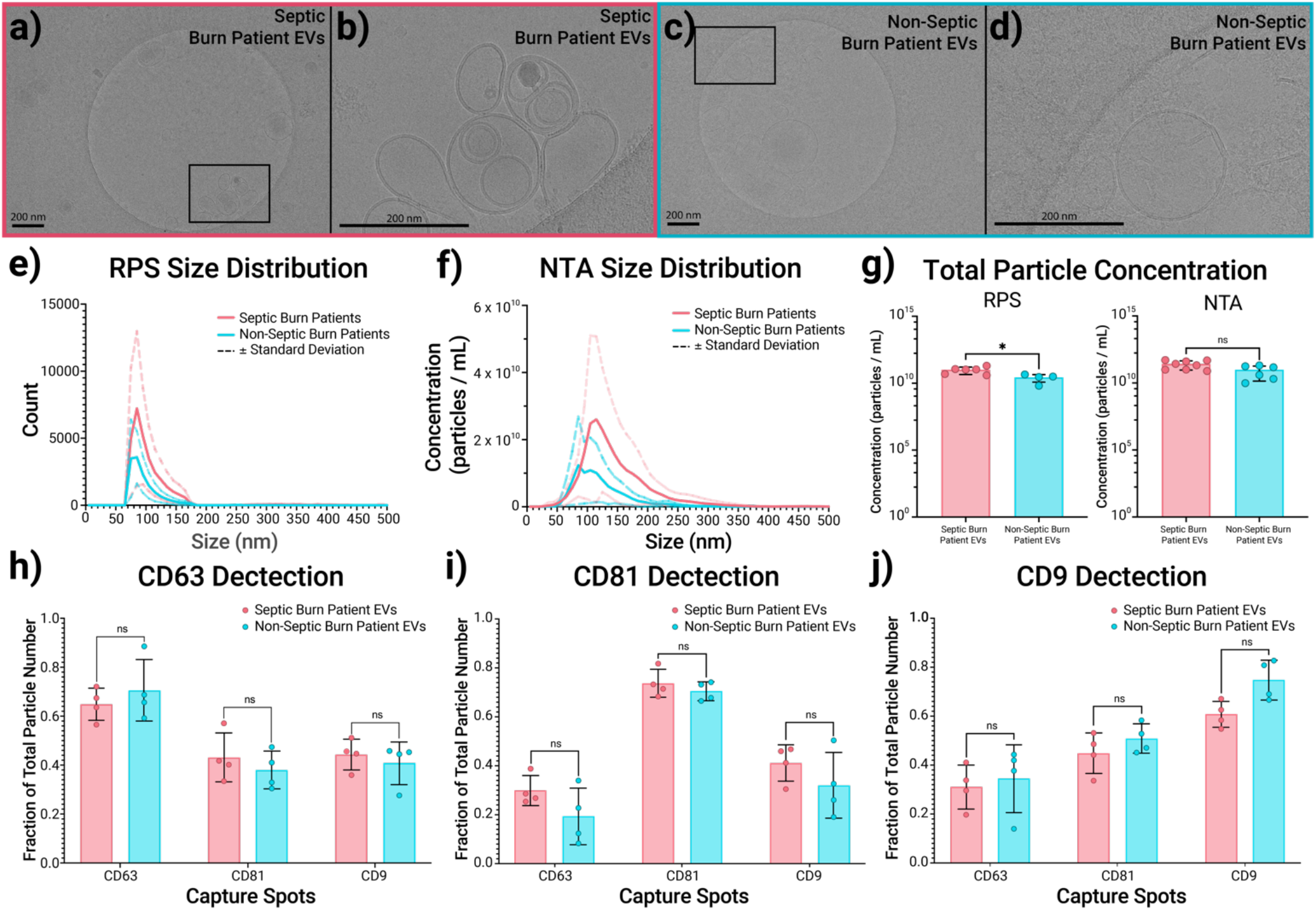
Characterization of SEC isolated EVs from septic and non-septic burn patient plasma. (a-d) CryoEM images of isolated EVs from (a, b) septic and (c, d) non-septic burn patient samples reveal the expected lipid bilayer structure. Images were acquired at (a, c) 11k and (b, d) 45k magnification. Scale bars are 200 nm. (e) RPS and (f) NTA were used to determine the average size distribution of the EVs isolated from the septic and non-septic burn patient samples. Dotted lines indicate intersample size standard deviation (g) The total EV particle concentration of the samples was measured by RPS and NTA. (h-j) Prescence and co-localization of human EV biomarkers (h) CD63, (i) CD81, and (j) CD9 determined by immunofluorescent tetraspanin kit assay.

Resistive pulse sensing (RPS) and nanoparticle tracking analysis (NTA) were used to determine the EV size distributions (**Figure 2e–f, Figure S1**). We found the average diameter of the septic burn patient EV samples to be 103.59 ± 0.28 nm (mean ± SE) and 141.46 ± 14.66 nm (mean ± SD) for RPS and NTA, respectively. This is compared the non-septic burn patient samples with an average diameter of 98.13 ± 0.28 nm (mean ± SE) and 130.07 ± 15.29 nm (mean ± SD) for RPS and NTA, respectively. Difference in average diameter between EVs isolated from septic and non-septic burn patient samples was found to be statistically significant for the NTA measurement (unpaired *t*-test, *p* = 0.0032). The NTA size distribution is a bell-shaped curve due to the limit of detection of EVs on NTA (∼ 80 nm) and the low light-scattering efficiency of EVs.^[44]^ On the other hand, the RPS size distribution shows a sharper peak near its respective limit of detection of ∼65 nm.

RPS and NTA were also utilized to determine the total concentration of particles isolated from the samples (**Figure 2g**). The average concentration of particles isolated from septic burn patient EV samples were 1.02*10^11^ and 2.68*10^11^ particles per mL for RPS and NTA, respectively. The average concentration for non-septic burn patient EV samples were 2.78*10^10^ and 9.90*10^10^ particles per mL for RPS and NTA, respectively. EVs isolated from the septic burn patient samples had statistically significantly higher concentration compared to those from the non-septic burn patients as determined by RPS (unpaired *t*-test, *p* = 0.0383). While a similar trend was seen for concentration measured by NTA, no statistically significant difference was found (p = 0.0523). It is possible that this increase in concentration between septic and non-septic burn patient EVs is related to the relative bacterial load of EVs between the two populations. Differences in concentrations between NTA and RPS measurements methods may be attributed to the presence of EVs smaller than the limit of detection of NTA versus RPS.^[44]^

The presence and co-localization of human EV biomarkers CD63, CD81, and CD9 in isolates was confirmed using an immunofluorescent tetraspanin kit assay (**Figure 2h–i, Figure S1**). No significant difference in tetraspanin expression was observed between the two EV populations (unpaired *t*-test, *p* > 0.05). Tetraspanin expression was measured with the immunofluorescent tetraspanin kit assays using microarrayed chips with antibody capture spots for human CD63, CD9, and CD81. It is of note that neither bacteria nor their EVs express these tetraspanins, although they do interact with them.^[45–47]^ Therefore, the tetraspanin kit assays would not capture bacterial EVs, and the observed difference in RPS measured concentration may be attributed to measuring additional EVs secreted from bacteria and/or due to patient secretion of EVs in response to infection. Guidance on bacterial EV specific markers or other characterization methods were not published at the time of data collection (e.g., MISEV 2018), and are limited to broad glycoconjugate markers like LTA and LPS for gram-positive and gram-negative bacterial classes respectively as recommendations from the updated MISEV 2023.^[42,43]^

### Raman Spectral Differences Between Septic and Non-Septic Burn Patient EVs

Following isolation and characterization of the EVs isolated from patient plasma, analysis via spontaneous Raman spectroscopy was performed. Burn patients were used as the control population in this study due to the overlapping clinical manifestations that burn patients and septic burn patients share in comparison to other patient populations (e.g., ill patients who have a topical bacterial infection that is not considered septic, or healthy controls). Comparison of the spontaneous Raman spectral profiles of EVs isolated from septic burn patients to non-septic burn patients represents a more realistic diagnostic scenario for this utilization case, where the standard diagnostic criteria fail to uphold.

Spontaneous Raman spectroscopic analysis was performed on the EV patient samples (n=8 septic burn patients and n=6 control non-septic burn patients). A 2 μL drop of each EV sample was deposited onto a quartz coverslip, dried, and analyzed on a custom-built inverted Raman microscope. A quick (90 s) spectral acquisition was performed for each sample at five different spots along the outer coffee-ring of the dried sample, where EVs tend to be enriched.^[48,49]^ Global spectral averages for septic versus control burn patients and their standard deviations are shown in **Figure 3a**. To investigate the differences between these patient population EV spectral profiles, we subtracted the spectral averages in MATLAB, where the difference between population averages = mean_septic_patients_ - mean_non-septic_patient_. The remnant spectrum of difference between population averages is shown **Figure 3b**. Positive peak intensities correlate to a higher normalized intensity of that peak in the septic burn patient EV average Raman spectrum, whereas negative intensities indicate a higher normalized intensity of that peak in the non-septic burn patient EV average spectrum.

**Figure 3:**
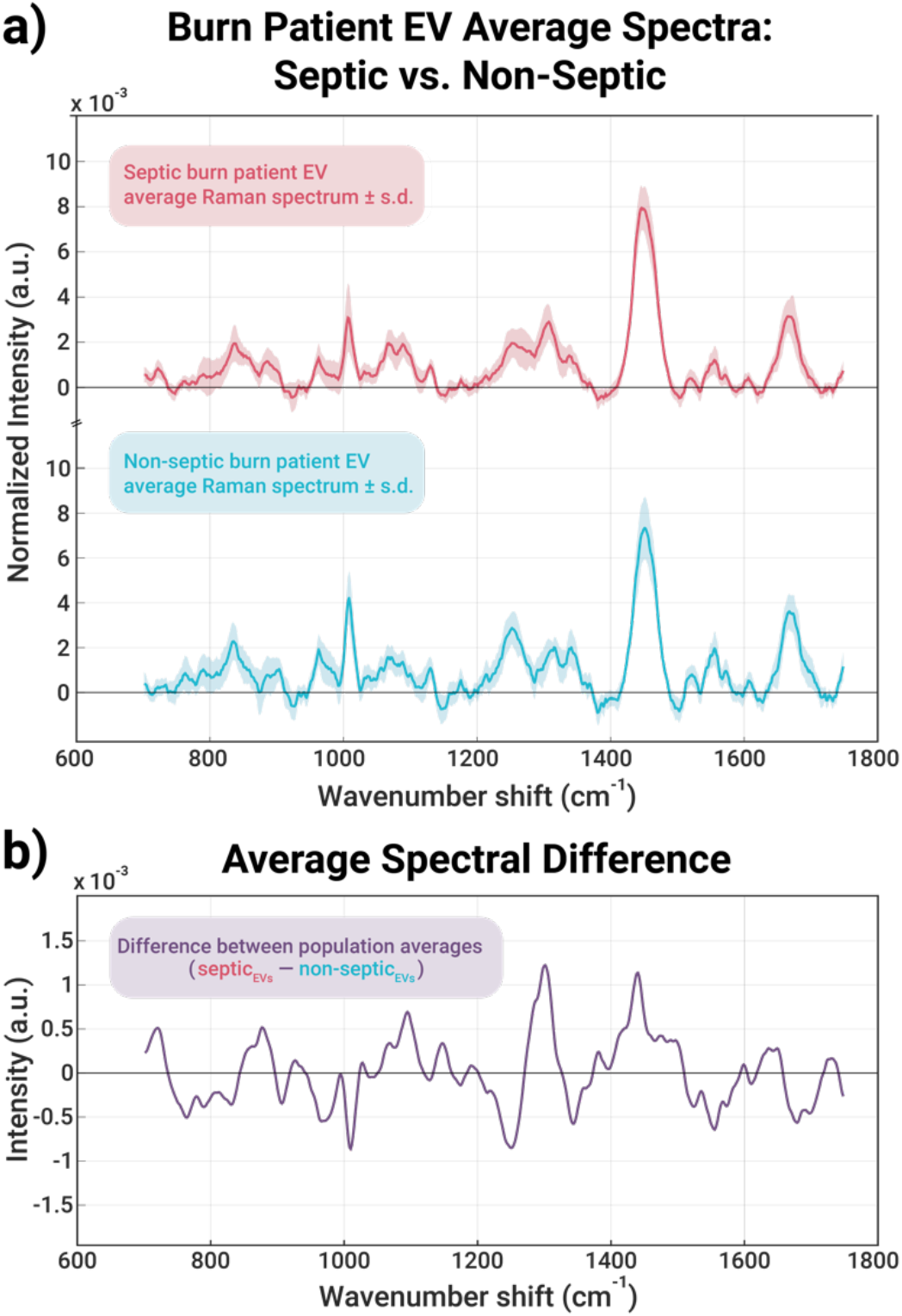
Global averages of septic and non-septic burn patient EVs. (a) Normalized global average and standard deviation spontaneous Raman spectra for septic burn patient EVs (n=8) (a), and for control non-septic burn patient EVs (n=6). For each patient plasma-isolated EV sample, a 90s spectrum was acquired at five different spots. (b) The normalized spectral difference between septic and non-septic burn patients Raman global average spectra.

Notable spectral differences include: an increased peak intensity at 718 cm^-1^ associated with CN^+^(CH_3_)_3_ stretching in the head group of phosphatidylcholine, an increased peak intensity at 877 cm^-1^ associated with C–C–N^+^ symmetric stretching in lipids and C–O–C ring vibrations in carbohydrates, a decreased peak intensity at 1005 cm^-1^ associated with the symmetric ring breathing of phenylalanine in proteins, increased peak intensities at 1068 cm^-1^ and 1095 cm^-1^, associated with PO_2_^-^ stretching in DNA and RNA, a decreased peak intensity at 1253 cm^-1^ in the Amide III region in the secondary structures of proteins, an increased peak intensity at 1301 cm^-1^ associated with the CH_2_ twisting mode in lipids, an increase at 1380 cm^-1^ correlated to RNA hairpins, an increase in C–H vibrations at 1440 cm^-1^ associated with the C–H bonds found across biological molecules, decreased intensity at 1535 cm^-1^ attributed to asymmetric NO_2_ stretch in the presence of a benzene ring, and the presence of a slight increase at 1731 cm^-1^ resulting from carbonyl stretching vibrations.^[50–53]^

### Differences in EV spectral profiles enable discrimination via PCA/QDA

To evaluate the utility of spontaneous Raman spectra of patient EVs as a proof-of-concept clinical indicator of sepsis in burn patients, we performed principal component analysis (PCA) and multivariate discriminant analysis. Raman spectral data were cropped, smoothed, and normalized using established methods.^[54–57]^ Normalization and PCA permitted dimensionality reduction of our dataset and limits feature scale sensitivity.^[58]^ Spectral data stacks were created for both the spectra acquired from the septic burn patient EV samples and from the non-septic burn patient EV samples. **Figure 4a** visualizes the individual patient spectra plotted in 2D-PC space along PC1 and PC2, encompassing nearly half (49.2%) of the total variance of the dataset.

**Figure 4:**
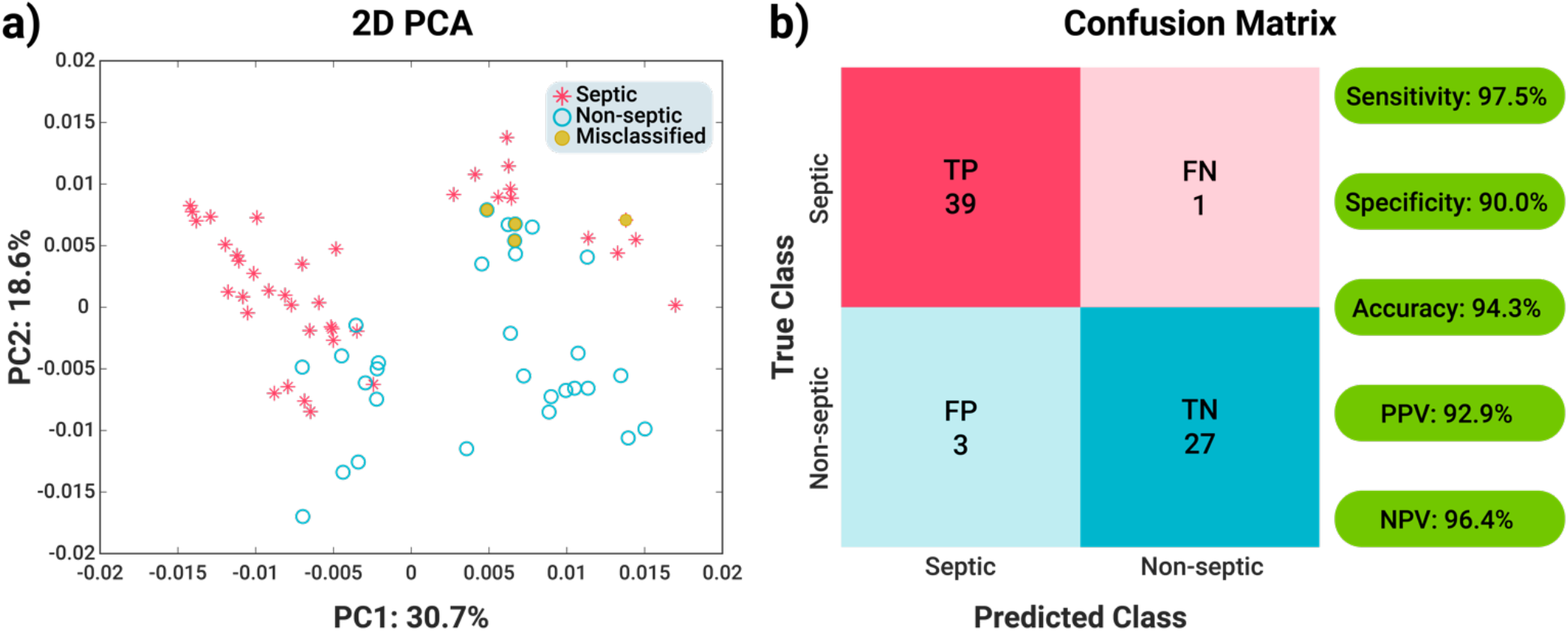
Principal Component Analysis and Confusion Matrix Results. (a) Normalized spontaneous Raman spectra acquired from patient EV samples (n=70) underwent principal component analysis and are plotted in PC space along PC1 and PC2. This 2D PC space encompasses 49.2% of the total variance of the dataset. Septic burn patient spectra are shown as pink stars, and non-septic control burn patients are labelled as blue circles. QDA analysis incorrectly identified four spectra, shown in a yellow overlay on their respective marker. Three spectra were found to be from the same non-septic burn patient (ARCS barcode number 1879), which were misclassified as septic (FP). One spectrum from a septic burn-patient was misclassified as non-septic (FN). (b) A confusion matrix visualizes the output of the optimized QDA analysis across the first five PCs (PC1 – PC5). Notably, just one of the 70 spectra was classified as a false negative result, yielding a high negative predictive value of 96.4% and a high sensitivity of 97.5%.

We performed supervised discriminant analytical data analysis to better visualize and model the separation of septic burn patients from non-septic (control) burn patients. Every combination of the first 5 principal components (PCs) (chosen as they represent >70% of the cumulative variance) were run through a custom MATLAB script to select the best data visualization model included within built-in MATLAB classifiers. PC loading coefficients are shown in **Supplemental Figure 2a–e**.

Quadratic discriminant analysis (QDA) of the first five PCs (PC1-PC5), accounting for 72.2% of the total cumulative percent variance, resulted in the highest combination of accuracy, sensitivity, and specificity of the patient clinical classifications (**Figure 4b)**. With this classifier, we have achieved 97.5% sensitivity and 90.0% specificity, with 94.3% accuracy in fitting the Raman spectra of the EV samples into PC space. Of the 70 patient spectra collected and passed through the classifier, just 1 was falsely predicted as non-septic and 3 were falsely predicted as septic, yielding a 2.5% false negative rate and a 10% false positive rate. A visual representation of the 70 measured spectra plotted in three-dimensional PC space (PCs 1–3), along with the quadradic hyperplane of best fit can be found in **Supplemental Figure 2f**.

While QDA analysis of the PC space caused a 10% incidence in false) positives (Type I error), it is notable that the false negative rate is low (2.5%) (Type II error. In this particular clinical indication, it is of the utmost importance to detect, then treat, septic burn patients as soon as possible. There are minimal effects of treating patients with antibiotics who ultimately do not prove to be septic (*i*.*e*., false positive), however it is critical that patients who are at risk of becoming, or already are, septic get the antimicrobial treatment, resources, and timely care they need to better their prognosis.^[59]^ Furthermore, sensitive, rapid identification of the bacteria present would help guide specific clinical treatment, lessening the need and time for systemic prophylactic antibiotics and potentially reducing the risk of downstream antibiotic resistance.^[59]^ In this case, the trade-off between sensitivity versus specificity of detection favors sensitivity. Thus, our high negative predictive value and low false negative rate support the premise of using spectral differences in the EV Raman fingerprint isolated from patient plasma samples.

For comparison, the use of procalcitonin (PCT) as a biomarker for sepsis in burn patients showed 82% sensitivity and 91% specificity following initial burn trauma but decreases to a combined 67% sensitivity and 87% specificity after this initial trauma phase, similar to the state of the patients in this study, as reported in a 2021 meta-analysis.^[60,61]^ Thus, our methodology shows promise for improving sensitivity and specificity when compared to current biomarkers of sepsis in burn patients.

Interestingly, three of the four misclassified spectra of the 70 acquired were from the same patient (ARCS patient barcode 1879), contributing to the Type I error, false positive. Spectral misclassifications could be interesting to examine with a larger data pool, to investigate other clinical indications or patient background information that could be driving the misclassifications. Exploring whether misclassifications correlate with specific patient comorbidities or treatment regimens could provide valuable information for personalized diagnostic approaches.

### Differences in patient EV Raman spectra are in part driven by the presence of bacterial EV glycoconjugate biomarkers

Common bacterial strains associated with septic bacteremia in both burn and non-burn patients include Gram-negative bacteria such as *Klebsiella* spp., *Escherichia coli*, and *Pseudomonas aeruginosa*, as well as Gram-positive organisms including *Staphylococcus aureus* and *Streptococcus* spp., among others.^[5,62,63]^ Since 1987, gram-positive bacteria have arisen as the predominant pathogens responsible for sepsis in the United States.^[64]^

We hypothesized that the spectral differences of septic versus non-septic burn patient EVs would be at least in part driven by the presence of pathogenic bacterial EVs shed from bacterial biofilm into the bloodstream. Thus, we sought to identify spectral markers of these bacteria-derived EVs to explain the Raman spectral differences of the EVs isolated from our two patient populations. The different biogenesis pathways of bacterially derived EVs cause them to have membrane glycoconjugate biomarkers

Briefly, bacterial EV (bEV) biogenesis is dependent on the group of bacteria they are derived from. Gram-negative bacterial EVs are commonly referred to as outer membrane vesicles (OMVs), and their biogenesis has been thought to be due to release from the outer membrane *via* reduction of outer membrane peptidoglycan–protein linkages, differential assembly of lipids/lipopolysaccharides (LPS) into microdomains, swelling pressure due to protein or peptidoglycan accumulation, or by explosive cell lysis.^[65]^ Gram-positive bEVs come about differently, due to the thick outer peptidoglycan layer of the bacterial cell cytoplasmic membrane complete with polyanionic matrix polymers such as macroamphiphiles like lipoteichoic acid (LTA), as well as the lack of the outer membrane structure that gram-negative bacterial cells possess.^[65,66]^ The different bEV biogenesis pathways effectively create vast heterogeneity within bEV/OMV types, however there are some distinguishing characteristics between them such as their bacterial glycoconjugates. Glycoconjugates are glycans with covalently linked species such as peptides, lipids, proteins, *etc*. We chose to investigate glycoconjugate markers for bacterial strains commonly associated with sepsis in burn patients. These chosen bacterial glycoconjugates and their origins are further described in **Table 1**.^[62,63,67–70]^

**Table 1:**
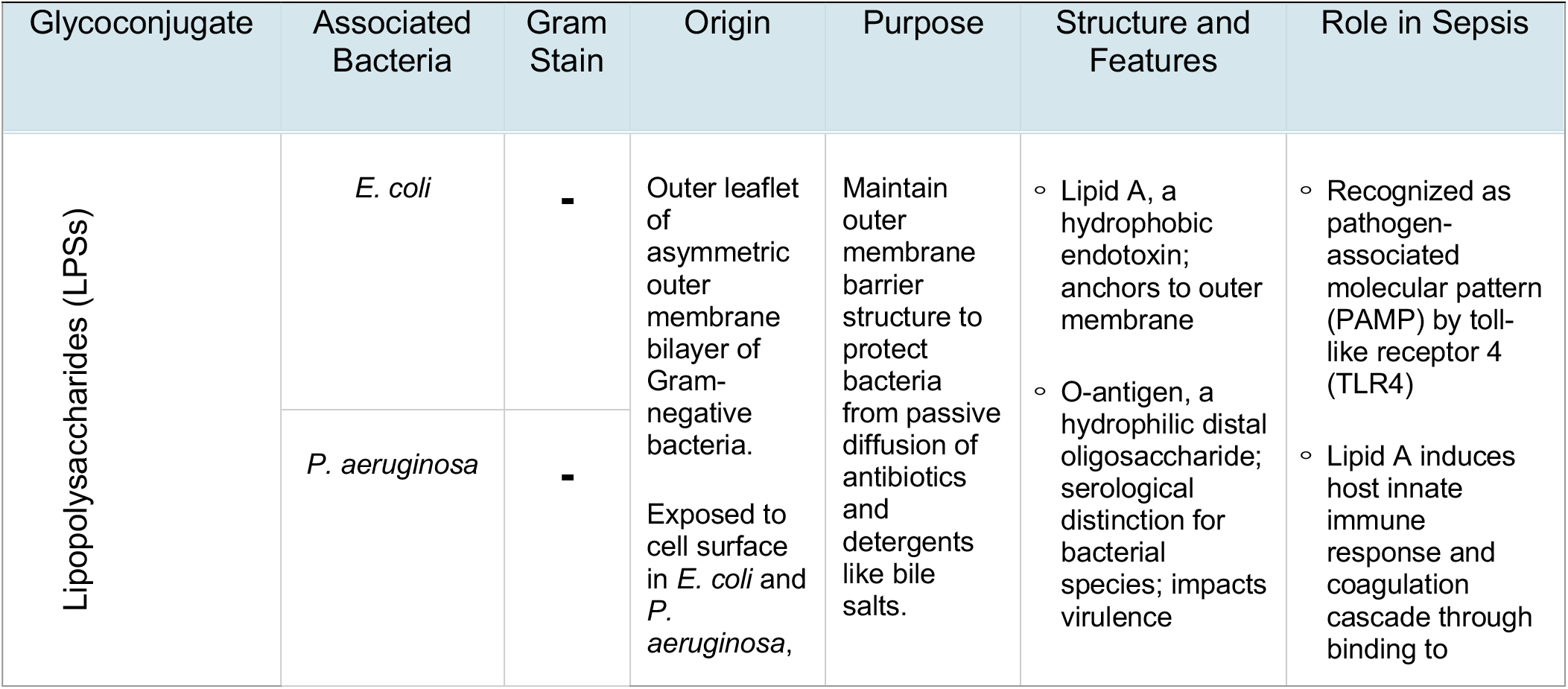
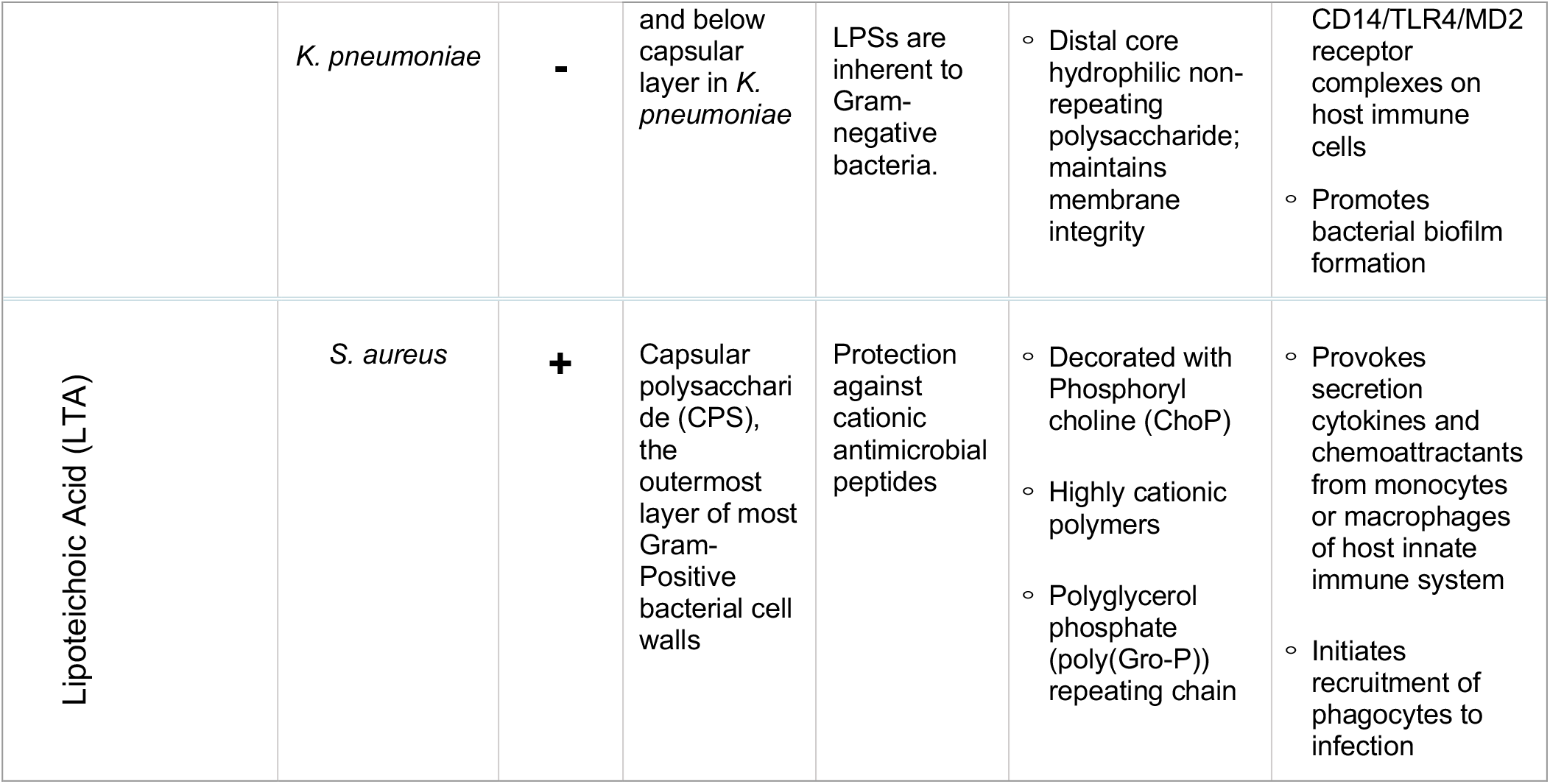
Characteristics of chosen bacterial glycoconjugate biomarkers.

It has been shown that discrete differences between enteric LPSs isolated from several gram-negative bacterial strains can be revealed through comparison of normalized surface-enhanced Raman scattering (SERS) spectra.^[71]^ Thus, to assess whether the spectral differences we noted in this study were at least in part driven by the presence of bacterial EV signatures, we measured lyophilized powder LPS analytical standards from *E. coli, K. pneumoniae, P. aeruginosa*, and an LTA analytical standard isolated from *S. aureus* (more information found in methods section). Raman spectra for each bacterial glycoconjugate standard were acquired with four replicates. These spectra were then normalized, smoothed, and background corrected for analysis.

To demonstrate that there are indeed unique spectral differences between these bacterial glycoconjugates that can be elucidated with spontaneous Raman, we performed PCA and hierarchical clustering. Clear spectral variance between each of these glycoconjugate standards was observed by their distinct hierarchical clustering in 3D PC space, as shown in **Figure 5a**. The generated average spectrum of the four replicates for each chosen glycoconjugate analytical standard after normalization, smoothing, and PLS fitting are shown in **Figure 5b**. After confirming discrete spectral differences between the glycoconjugate analytical standards, we compared them to the spectral difference of the population average spectra of septic and non-septic burn patient EVs (**Figure 3b**). The asymmetric least squares fit of these standards to the difference between population averages Raman spectrum is shown in **Figure 5c**.

**Figure 5:**
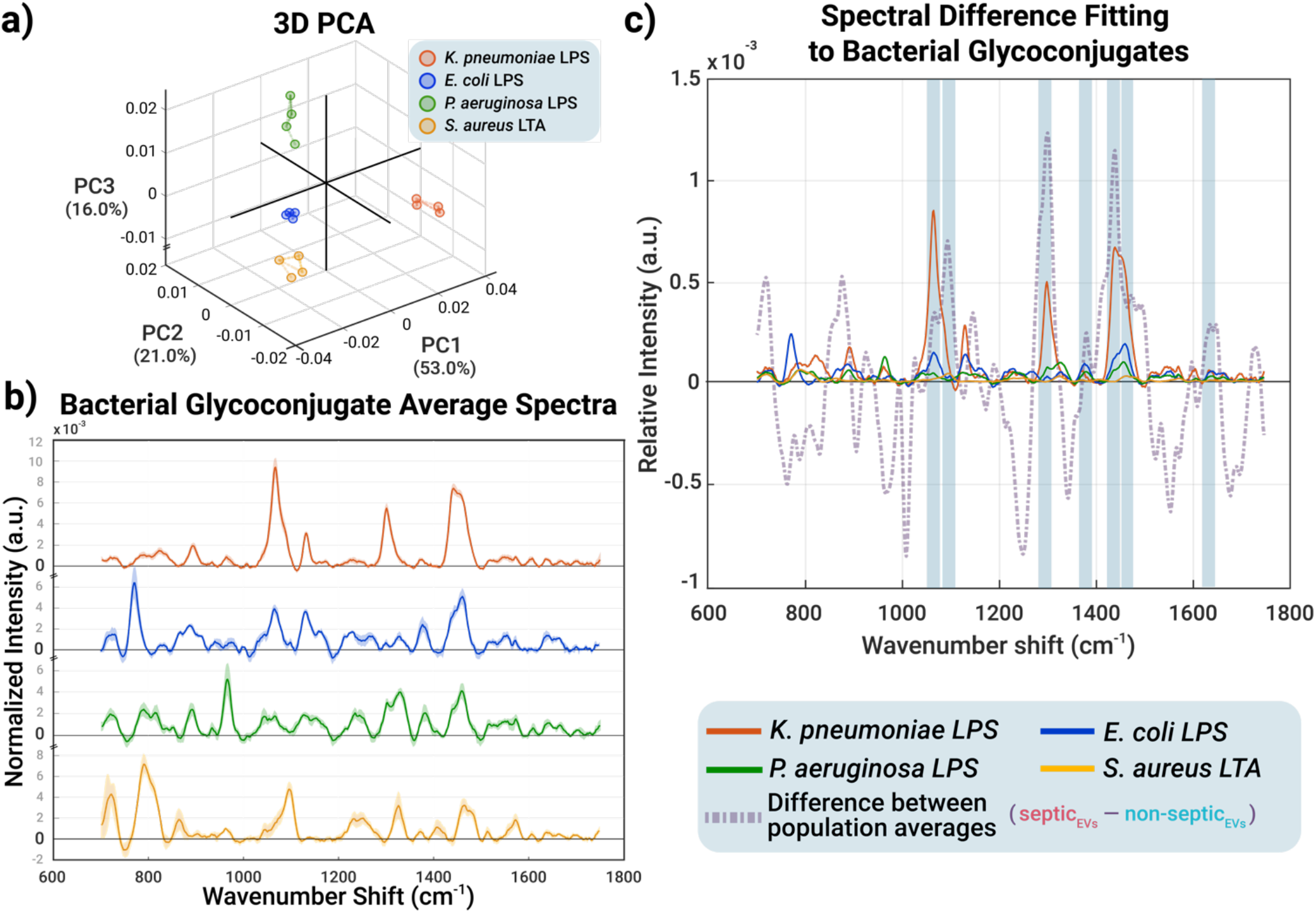
Utilizing bacterial glycoconjugates to elucidate spectral differences between septic and non-septic burn patient EV Raman fingerprints. (a) Clustering analysis of the four chosen bacterial glycoconjugates’ Raman spectra visualized in three-dimensional principal component space across the first three PCs, accounting for 90% of the variance. Clear clustering validates spectral differences between the four standards for downstream use as reference analytes. (b) The average spontaneous Raman spectrum of each chosen bacterial glycoconjugate analytical standard following normalization, smoothing, and PLS fitting. Each standard was measured four times then averaged. Shading shows the standard deviation. (c) Comparison of the difference of the average spectrum of non-septic burn patient EVs from septic burn patient EVs (denoted as difference between population averages, in purple) to the four chosen bacterial glycoconjugates of bacteria commonly associated with sepsis in burn patients. Raman band fits are highlighted in blue. Key in bottom right corner of figure is for panels (b) and (c).

While there are some clear feature overlaps with the sepsis-related EV spectral features to the bacterial glycoconjugate standards, there is not one fit that is overwhelmingly strong. Spectral peak fitting shows some overlap with lipopolysaccharides isolated from *K*. pneumoniae, *E. coli, P. aeruginosa*, respectively. Fit with *S. aureus* LTA was only observed at 1095 cm^-1^. These results are summarized in **Table 2**.^[70–73]^ It is understandable that the total variance between the Raman signatures of the two EV populations would not be fully explained by these four specific bacterial glycoconjugates.^[43]^

**Table 2:** Fits for Raman spectra of bacterial glycoconjugates to the difference of population average spectrum.

| Raman Band | Vibrational Mode and Assignment | Associated Glycoconjugate(s) |
| --- | --- | --- |
| 1067 $\text{cm}^{-1}$ | C–C and C–O stretching vibrations of O-Antigen and distal LPS core<sup>[77]</sup> | *K. pneumoniae* LPS<br>*E. coli* LPS |
| 1095 $\text{cm}^{-1}$ | P–O stretching of poly(Gro-P) chain<sup>[70]</sup> | *S. aureus* LTA |
| 1301 $\text{cm}^{-1}$ | C–H deformation and CH<sub>2</sub> bending of O-Antigen and distal LPS core<sup>[77]</sup> | *K. pneumoniae* LPS<br>*E. coli* LPS<br>*P. aeruginosa* LPS |
| 1380 $\text{cm}^{-1}$ | CH<sub>3</sub> deformation and C–N stretching in O-Antigen<sup>[72]</sup> | *K. pneumoniae* LPS<br>*E. coli* LPS |
| 1440 $\text{cm}^{-1}$ | CH<sub>2</sub> scissoring in Lipid A<sup>[72]</sup> | *K. pneumoniae* LPS<br>*E. coli* LPS<br>*P. aeruginosa* LPS |
| 1455 $\text{cm}^{-1}$ | CH<sub>2</sub>/CH<sub>3</sub> scissoring, C–O–C deformation in glycosidic linkages, C–C–C deformation, and C–O–H deformation in Lipid A<sup>[72]</sup> | *P. aeruginosa* LPS |
| 1637 $\text{cm}^{-1}$ | C–O and C–C stretching within Lipid A<sup>[72]</sup> | *K. pneumoniae* LPS<br>*E. coli* LPS |

Other work has investigated SERS spectral classification of bacterial EVs from *E. coli* strains, as well as SERS for six typical bacteria contributing to nosocomial infections, however, there has yet to be a study into spontaneous Raman of bacterial EVs isolated from patient biofluids.^[41,39]^ Differentiation of systemic inflammatory response syndrome and sepsis from plasma using Raman spectroscopy has been reported, but was not specific to sepsis in burn patients and did not investigate the roles of bacteria, EVs, or bacterial EV signatures.^[74]^

The field of bacterial EV characterization still requires further advancement. Specific biomarkers for bacteria-derived EVs have yet to be identified and agreed upon, and RS-based EV characterization is still in its infancy, thus large datasets for comparison across studies are not available.^[43]^ Differential underlying patient microbiota, or different pathogenic bacterial strains may also be a driving contribution to the spectral separation and are fields of interest for future study.^[75,76]^

Ultimately, while these results are promising, limitations of our work include a low patient number due to the niche patient population, precluding more advanced deep learning to improve model performance. The relatively small sample size of the study is further attributed to the use of remnant plasma samples derived from routine clinical chemistry samples drawn at regular intervals, rather than at the time of initial sepsis recognition. Further prospective collection of samples and associated clinical course will build on this work. Other potential limitations in the burn population are the aforementioned difficulties in identification of sepsis, as burn patients exhibit the inflammatory response syndrome throughout their hospital course. Burn patients may also exhibit transient bacteremia from dressing changes and/or surgical excision of heavily colonized wounds. An early marker that distinguishes these different states, such as the EV-Raman workflow described in this work, will help to move the science of sepsis forward.

Nevertheless, we recognize the utility and translatability of RS analysis of EVs for detection and diagnosis of other bacterial bloodstream infections. For example, staph infections, caused by gram-negative bacteria *Staphylococcus aureus*, have a mortality rate around 30%.^[78]^ Current diagnostic methods rely on culture-based methods spanning multiple days, or genomic/proteomic methods which require highly specialized equipment and technicians.^[79,80]^ Earlier diagnosis of staph infection has been shown to decrease relative mortality by 75.5%.^[81]^ Therefore, employing a similar workflow to that described in this work for early diagnosis of staph infection *via* Raman could be used decrease the mortality of patients. Raman EV analysis could also be translated to diagnosis other infectious diseases caused by bacteria. 1.6 million people died of tuberculosis (TB) in 2021, caused by *M. tuberculosis*. Efficient diagnosis of TB is critical for effective treatment of TB, which is estimated to have saved 74 million lives from 2000 to 2021.^[82,83]^ EVs secreted by *M. tuberculosis* bacteria have been established as a reliable biomarker of TB and therefore Raman EV analysis following our workflow may be a useful for diagnosis of TB.^[84]^

## Conclusion

The focus of this work was to utilize Raman spectroscopy to spectrally discriminate markers of sepsis in EVs isolated from burn patient plasma. We show, for the first time, the possibility of utilizing spontaneous Raman scattering and resultant spectra of patient plasma isolated extracellular vesicles for the detection of sepsis in burn patients. Through quadratic discriminant analysis of the acquired patient EV Raman spectra, we were able to separate septic from non-septic burn patient EVs with 97.5% specificity and 90.0% sensitivity, and in part attribute the spectral differences to the presence of bacterial EV markers of four common bacterial strains that contribute to sepsis morbidity such as lipopolysaccharides for *K. pneumoniae, E. coli*, and *P. aeruginosa* and lipoteichoic acid isolated from *S. aureus*. Overall, we contributed to the discourse surrounding bEVs and OMVs for pathogenic detection, but there is still much work to be explored in this field regarding their isolation and characterization. Collectively, our proof-of-concept findings establish a workflow for future clinical applications of spectral investigation into bacterial EVs within host biofluids and show the promise of Raman spectroscopy of patient plasma isolated EVs for early identification and stratification of sepsis in vulnerable burn patient populations.

## Methods

### Patient Plasma Sample Collection

Both control (non-septic) and septic patient plasma samples were provided by the University of California, Davis Pathology Biorepository. Samples were derived from residual clinical chemistry plasma samples collected from adult burn patients at the burn intensive care unit (BICU) at the University of California, Davis Health Fire Fighters Burn Institute Regional Burn Center. Samples were collected with accordance to the UC Davis Institutional Review Board (IRB #1911992) and utilized the UC Davis Pathology Biorepository anonymous clinical research specimen protocol. Under the ARCS, only patient demographics such as age, gender, race/ethnicity, and quality of specimen, collection container, etc. is available to the researcher. Patient data is stripped of identifiers and assigned an ARCS barcode number. Available patient data is summarized in **Supplemental Table 1**.

### EV Isolation

All work with EVs is performed utilizing Protein Lo-Bind Eppendorf tubes (Catalog No. 022431081, Eppendorf; DE), and Lo-Bind pipette tips (Corning DeckWorks, Corning Inc.; NY, USA). Plasma from control and sepsis patients was centrifuged in Protein Lo-Bind Eppendorf tubes (Catalog No. 022431081, Eppendorf; DE) at 5,000 *x g* for 10 min to remove cells and cell debris. The supernatant was collected and stored at -80°C until further isolation. Izon qEV single 35 nm SEC columns and automatic fraction collector (AFC) v2 with firmware version V1.3.1 were then used to isolate the EVs (Izon Science Ltd; NZ). The storage buffer was removed and 6 mL of 0.2 μm filtered PBS was used to flush the columns. 100 μL of sample was added to the column followed by 3 mL of 0.2 μm filtered PBS after the sample moved through the column frit. If samples needed to be diluted to reach this 100 μL volume, they were diluted in 0.2 μm filtered PBS. A single fraction of 0.8 mL was collected after collecting an initial buffer volume of 0.87 mL. The collected fraction was aliquoted to prevent freeze-thaw cycling and stored at -80°C until further analysis was performed. Columns were not re-used.

### CryoEM

Copper girds (Quantifoil, 1.2 μm diameter holes, 1.3 μm separation, 300M Cu grid) were glow discharged for 25 seconds at 15 mA negative polarity (PELCO easiGlow, Ted Pella Inc.; CA, USA). The grids were then loaded onto the Leica EM GP2 Plunge Freezer (Leica Microsystems Inc., IL, USA). The chamber was kept at 18°C and 80% humidity, and the cryogen was kept at -180°C. 4 μL of sample was then incubated onto the grid for 2 minutes, blotted for 6 seconds, then flash frozen. Grids were stored in liquid nitrogen until imaged. Imaging of the grids was performed on a FEI Talos L120C Transmission Electron Microscope (Thermo Fisher Scientific Field Electron and Ion Company (FEI); OR, USA). Images were analyzed using Adobe Photoshop 2023 (version 24.4.1, Adobe; CA, USA) and contrast was increased to 100.

### Nanoparticle Tracking Analysis (NTA)

Nanoparticle Tracking Analysis (NTA) was performed using a NanoSight model LM10 (Malvern Panalytical Ltd.; UK), equipped with a 405 nm laser and sCMOS camera. SEC-isolated EV samples were thawed at room temperature from aliquots stored at -80°C. Samples were diluted 40-240x in 0.02 μm filtered PBS to not oversaturate the detector. Prior to data acquisition, NTA lines were flushed first with 70% ethanol then with 0.02 μm filtered PBS at 4°C to ensure no background signal prior to sample addition. Typically, 1 mL of sample is loaded into a 1 mL syringe and placed within an automated syringe pump (Harvard Bioscience; MA, USA). Three 30 second video acquisitions were recorded at camera level 12 for each sample. NanoSight NTA 3.1 software was used with detection threshold 2 and default screen gain 10 to track EVs while minimizing any background artifacts. Results for averaged sepsis and control samples shown in **Fig. 2f-g**. Size distribution graph for each sample is provided in **Fig. S1**.

### Resistive Pulse Sensing (RPS)

Resistive pulse sensing (RPS) was performed on nCS1 (Spectradyne; CA, USA) using C-400 cartridges (Spectradyne; CA, USA). Buffer solution of 1% Tween 20 and 1% PBS filtered with 0.2 μm non-pyrogenic filter was freshly prepared prior to each use to lower the surface tension of water. To prime the instrument, a reusable cleaning cartridge was run. Internal standard diluting 200 nm bead (3000 Series Nanosphere Size Standard, Thermo Fisher Scientific; MA, USA) solution was prepared by diluting bead solution by 1000 times in buffer solution that was further filtered by a 0.02 μm filter. EV samples were then diluted in the bead solution for a final concentration between 6*10^9^ and 2*10^10^ particles per mL (minimum of 1:1 dilution). 3 μL of the EV and bead solution was run on the C-400 cartridge. RPSpass software (version 1.0.2, National Cancer Institute; MD, USA) was then used analyze the data and remove outliers and bead particles. Results for averaged sepsis and control samples shown in **Fig. 2e, g**. Size distribution graph for each sample is provided in Fig. S1.

### Single-Particle Interferometric Reflectance Imaging Sensing (SP-IRIS)

Leprechaun Exosome Human Tetraspanin Kits (Cat. 251-1044, Unchained Labs; CA, USA) were stored in 4°C and chips were warmed to room temperature prior to use. Chips were pre-scanned to identify any pre-existing adhered particles according to the provided protocol. Isolated EV samples were diluted in 1x incubation solution (provided 10x incubation solution was diluted to 1x concentration with Ultrapure Type I water (18.2 MΩ·cm, PURELAB Chorus, ELGA LabWater, Veoilia Water UK Limited; UK) with 0.2% BSA) to achieve between 1000 and 6000 particles per capture spot. For sample incubation, chips were placed in the provided 24-well plate and 35 μL of diluted EV sample was pipetted onto the center of the chip. The 24-well plate was then sealed, covered in aluminum foil, and allowed to incubate overnight. After incubation, 300 μL per chip of 1x blocking solution and 0.6 μL per chip of anti-CD9, anti-CD63, and anti-CD81 antibody was prepared. Chips were then washed with the ExoView CW1000 Plate Washer (Unchained Labs; CA, USA) on the CW-TETRA protocol. 250 μL per chip of prepared blocking solution was added when prompted. Chips were then transferred to the chuck and imaged using ExoView R100 Automated Imager (Unchained Labs; CA, USA). Data was analyzed using ExoView Analysis Software (Unchained Labs; CA, USA). Fluorescent cutoffs were chosen to limit the particle count on the MIgG capture spot to be approximately less than 100 events and used for all chips (red channel: 200 a.u., green channel: 250 a.u., blue channel: 450 a.u.). Results for averaged septic and non-septic samples shown in **Fig. 2h-j**. Tetraspanin expression for each sample is provided in **Fig. S1**.

### Raman spectral acquisition of EV samples

SEC-isolated EV samples were thawed at room temperature from aliquots stored at -80°C. Each sample was homogenized with a Lo-Bind pipette tip, and a 2 μL drop of each sample was placed on 25 mm round quartz coverslips, thickness #1 (0.15-0.18 mm) (SPI Supplies; PA, USA). Note that control patient samples 1880 and 1890 required a second 2 μL droplet to dry to increase the concentration of EVs within the sample for better signal acquisition. Spots were allowed to dry on a hotplate at 45°C for 5 minutes prior to measurement. All Raman spectra were acquired using our custom-built inverted Raman scanning confocal microscope with an excitation wavelength of 785 nm. Sample focus was performed manually under a 60x 1.2 NA water immersion objective on an inverted Olympus IX73 microscope (Olympus Life Science; MA, USA). Spectral capture was performed using an Andor Kymera 328i-C spectrophotometer (Andor Technology Ltd, Oxford Instruments; UK), and both spectral and brightfield images were captured using a Newton DU920P-BR-DD CCD camera (Andor Technology Ltd, Oxford Instruments; UK). Brightfield images were collected using a 0.30 NA lens. Andor Solis version 4.30.30034.0 software (Andor Technology Ltd, Oxford Instruments; UK) was used for acquisition and initial processing. Prior to each day of measurements, the instrument was calibrated for both wavenumber and intensity, using a Neon reference lamp and NIST traceable SL1-CAL VIS-NIR Tungsten Halogen lamp (StellarNet Inc.; FL, USA), respectively. Five spectra (90 second exposure) were sampled from each patient sample (25 mW laser power at sample, 785 nm excitation wavelength). Spectral data was acquired with a center wavelength of 850 nm to avoid laser interference, 2 mm aperture, and with a grating of 3 (600 l/mm, blaze 750 nm), allowing for collection of spectra from 789.42 nm to 909.88 nm (71.33 cm^-1^ – 1748.39 cm^-1^). Spectra containing cosmic rays were not included.

### Raman spectral acquisition of LPS and LTA Standards

Lipopolysaccharides from *Escherichia coli* O55:B5 irradiated (Product Number L6529), Lipopolysaccharides from *Klebsiella pneumoniae* - purified by phenol extraction (Product Number L4268), Lipopolysaccharides from *Pseudomonas aeruginosa* 10.22 source strain ATCC 27316. - purified by phenol extraction (Product Number L9143) and Lipoteichoic acid from *Staphylococcus aureus* (Product Number L2515) were purchased from Sigma Aldrich. Glycoconjugate standards were diluted in Ultrapure Type I water (18.2 MΩ·cm, PURELAB Chorus, ELGA LabWater, Veoilia Water UK Limited; UK) based on their respective solubilities, then 2 μL of each glycoconjugate standard was dried on 25 mm round quartz coverslips, thickness #1 (0.15-0.18 mm) (SPI Supplies; PA, USA) on a hotplate at 45 °C prior to RS acquisition. LPS from E. coli was diluted to 6 mg/mL, LPS from K. pneumoniae was diluted to 4 mg/mL, LPS from P. aeruginosa was diluted to 6.6 mg/mL and LTA from S. aureus was diluted to 2.2 mg/mL. Each sample underwent Raman spectral acquisition four times. Acquisition parameters and setup were otherwise the same as for aforementioned patient EV samples. Following spectral processing, an average spectrum for each standard was made by averaging the four acquisitions.

### Raman Spectral Processing

Spectral processing was performed using an in-house spectral processing GUI script built with MATLAB and carried out using MATLAB r2020b (MathWorks; MA, USA). All spectra were cropped from 700 cm^-1^ to 1800 cm^-1^, background corrected with penalized least-squares, smoothed with a Whitaker Smoothing filter, and intensity normalized using trapezoidal numerical integration so the AUC = 1 for spectral comparisons. Subsequent spectral analysis including principal component analysis, clustering, linear and quadratic discriminant analysis were also performed utilizing custom-built MATLAB scripts and built-in MATLAB functions/toolboxes.

### Statistical Analysis

RPS and NTA average size distribution for sepsis and control samples were determined using GraphPad Prism 10 (Version 10.1.0; GraphPad; San Diego, CA, USA) and Microsoft Excel (Microsoft; Redmond, WA, USA). Values shown for particle total concentration (RPS and NTA) and tetraspanin expression are the indicated as the mean ± SD. Statistical analysis for particle total concentration (RPS and NTA) was determined using unpaired *t*-test assuming Gaussian distribution with GraphPad Prism 10 (Version 10.1.0; GraphPad; San Diego, CA, USA)). Statistical analysis for tetraspanin expression were determined using unpaired *t*-test assuming Gaussian distribution with GraphPad Prism 10 (Version 10.1.0; GraphPad; San Diego, CA, USA). All experiments were performed with replicates; NTA and RPS was performed with three replicate runs per sample, multiple CryoEM grids were prepared for representative samples for each group, and each Raman measurement was performed five times for each patient sample and four times for the glycoconjugate standards. Statistical significance was considered when p< 0.05.

## Author Contributions

H.J.O and R.P.C. conceptualized and designed the study. H.J.O. performed the bulk of the data acquisition and analysis and drafted the manuscript. N.L contributed to EV sample isolation, characterization, and analysis, and drafting of the manuscript. V.A. contributed EV characterization. A.V.K. contributed Raman spectral acquisition. N.K.T. and T.L.P. revised the manuscript. N.K.T. contributed to the study design and provided clinical samples. R.P.C. developed the software utilized in the work, revised the manuscript, and obtained research funding supporting the project.

## Acknowledgments

Components of figures in this work were created using BioRender.com. Specimens were provided by the UC Davis Pathology Biorepository. Cryo-electron microscopy was performed at the UC Davis BioEM Facility in Briggs Hall.

