## Supplemental material for "Plasma-derived Extracellular Vesicles (EVs) as Biomarkers of Sepsis in Burn Patients via Label-free Raman Spectroscopy"

### Supplemental Table 1: De-identified Patient Information

| ARCS Patient Barcode | Sepsis status at time of blood draw consent | Sepsis status at time of experiments | Age (years) | Sex (M/F) | PCT [ng/mL] | Total Body Surface Area of Burn (TBSA) (%) |
|---|---|---|---|---|---|---|
| 1777 | Maybe | Yes | 37 | M | 6.22 | 51 |
| 1779 | Maybe | Yes | 47 | M | 3.95 | 33 |
| 1781 | Yes | Yes | 63 | M | 91.4 | 45 |
| 1782 | Yes | Yes | 54 | F | 9.3 | 25 |
| 1785 | Yes | Yes | 38 | M | 1.92 | 21 |
| 1876 | Yes | Yes | 32 | M | 21.5 | 42.5 |
| 1883 | Yes | Yes | 26 | M | 12.4 | 39.5 |
| 1884 | Yes | Yes | 41 | M | 9.63 | 21.5 |
| 1877 | No | No | 44 | F | 4.2 | 28 |
| 1878 | No | No | 28 | F | 3.68 | 19 |
| 1879 | No | No | 32 | M | 3.18 | 17 |
| 1880 | No | No | 59 | M | 2.22 | 35 |
| 1890 | No | No | 36 | M | 3.57 | 18.5 |
| 1891 | No | No | 59 | F | 2.49 | 29 |

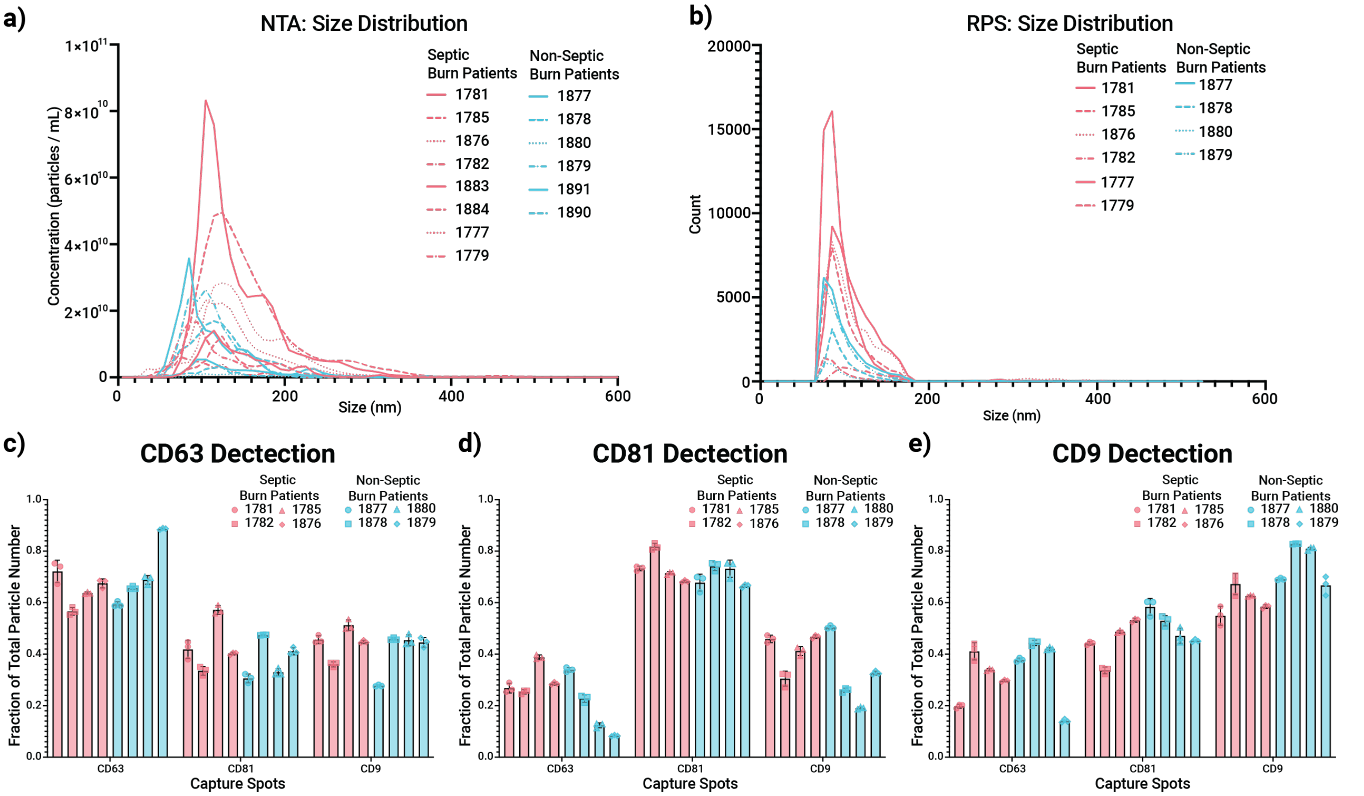


**Supplemental Figure 1: Characterization of SEC-isolated EVs from septic and non-septic burn patients.** (a) NTA and (b) RPS were used to determine the size distribution of the EVs. (c-e) Prescence and co-localization of mammalian EV biomarkers (c) CD63, (d) CD81, and (e) CD9, as determined by immunofluorescent tetraspanin kit assays.

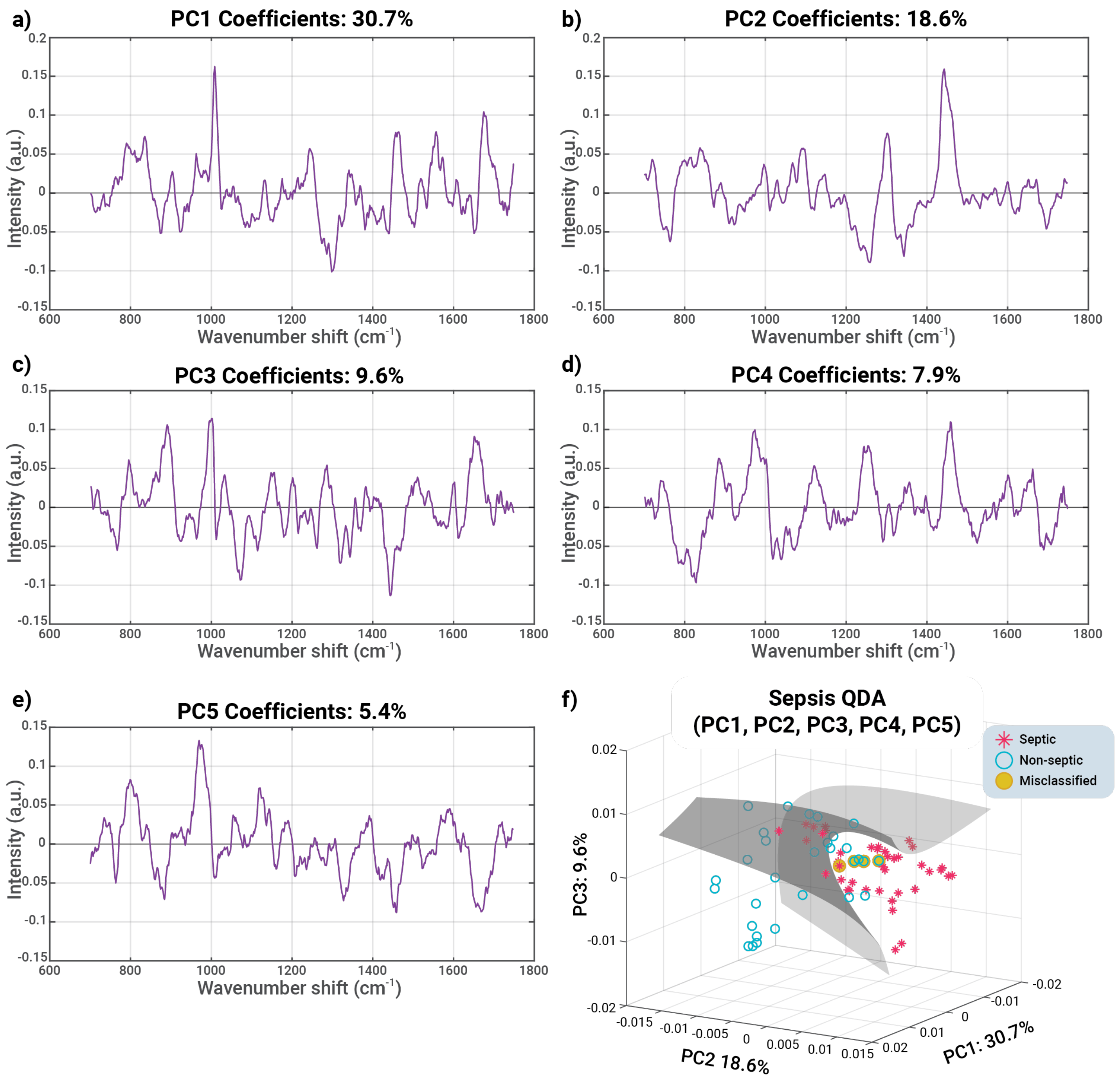


**Supplemental Figure 2: Principal Component Analysis Individual PC Loadings.** (a-e) Spectra of PC loadings for PCs 1 – 5. (a) PC1 accounted for 30.7% of the variance. (b) PC2 accounted for 18.6% of the variance. (c) PC3 accounted for 9.6% of the variance. (d) PC4 accounted for 7.9% of the variance. (e) PC5 accounted for 5.4% of the variance. These first 5 PCs accounted for a total of 72.2% of the variance of the spectra when comparing EVs isolated from septic burn patient plasma versus non-septic burn patient plasma. (f) Three-dimensional quadratic discriminant analysis of PCs 1 – 5 visualizes a quadratic hyperplane of best separation, showing separation of septic burn patient EVs (pink stars) from non-septic burn patient EVs (blue circles), and misclassified patient EV spectra overlayed with yellow circles.
